# Genome-wide association and prediction reveals the genetic architecture of cassava mosaic disease resistance and prospects for rapid genetic improvement

**DOI:** 10.1101/031179

**Authors:** Marnin D. Wolfe, Ismail Y. Rabbi, Chiedozie Egesi, Martha Hamblin, Robert Kawuki, Peter Kulakow, Roberto Lozano, Dunia Pino Del Carpio, Punna Ramu, Jean-Luc Jannink

## Abstract

Cassava (*Manihot esculenta*) is a crucial, under-researched crop feeding millions worldwide, especially in Africa. Cassava mosaic disease (CMD) has plagued production in Africa for over a century. Bi-parental mapping studies suggest primarily a single major gene mediates resistance. To be certain and to potentially identify new loci we conducted the first genome-wide association mapping study in cassava with 6128 African breeding lines. We also assessed the accuracy of genomic selection to improve CMD resistance. We found a single region on chromosome 8 accounts for most resistance but also identified 13 small effect regions. We found evidence that two epistatic loci and/or alternatively multiple resistance alleles exist at major QTL. We identified two peroxidases and one thioredoxin as candidate genes. Genomic prediction of additive and total genetic merit was accurate for CMD and will be effective both for selecting parents and identifying highly resistant clones as varieties.

---

Cassava (*Manihot esculenta* Crantz) is a crucial staple food crop, usually grown by smallholder farmers and feeding over half a billion people worldwide, especially in sub-Saharan Africa (http://faostat.fao.org). Breeding cycles are long in this outcrossing, clonally-propagated crop and genetic gains from breeding have been small over the last century compared with other crops (Ceballos et al., 2004, 2012). With a recently sequenced genome (Prochnik et al., 2012) and a high-density SNP-based genetic linkage maps ((ICGMC), 2014), it is for the first time possible to study the genetic architecture of key traits using modern genome-wide association analysis (GWAS) and to improve those traits with genomic selection (GS) (Oliveira et al., 2012; Ly et al., 2013).

Cassava mosaic disease (CMD) is the longest running and thus-far most impactful of the challenges cassava farmers face in sub-Saharan Africa(Fauquet et al., 1990). The disease is caused by several related species of geminiviruses and transmitted both through infected cuttings and by a vector, the common whitefly (*Bemisia tabaci* G.). Development and deployment of resistant cultivars is the most effective control method for this devastating disease. Following an unsuccessful world-wide search for resistance in cultivated germplasm (*M. esculenta*) in the 1930s, cassava breeders at the Amani research station in Tanzania resorted to interspecific hybridization with Ceara rubber tree (*M. glaziovii* Müll. Arg) and other related wild species in the 1930s (Hahn et al., 1979, 1980a; Fauquet et al., 1990). Moderate polygenic resistance combined with reasonable root yields was achieved through several cycles of backcross of Ceara rubber to the cultivated cassava (Hahn et al., 1980b). One of these interspecific hybrids, clone 58308, was subsequently used to initiate cassava breeding breeding at the International Institute of Tropical Agriculture (IITA) in the 1970s and resulted in the Tropical Manihot Selections (TMS) varieties (Hahn et al., 1980b).

More recently, a strong qualitative and dominant monogenic resistance known as *CMD2* was discovered in a Nigerian landrace (TMEB3) in the 1980’s (Akano et al., 2002). Multiple bi-parental QTL analyses have been conducted, initially using SSR markers (Akano et al., 2002; Lokko et al., 2005; Okogbenin et al., 2007, 2012a; Mohan et al., 2013) but more recently genome-wide SNPs (Rabbi et al., 2014a; b) to understand the genetic basis of this type of qualitative resistance to CMD. Although some studies hint at additional resistance loci (Okogbenin et al., 2012a; Mohan et al., 2013) most evidence points solely to the *CMD2* locus (Rabbi et al., 2014a; b). However, these bi-parental mapping efforts relied on a handful of unique parental genotypes from West Africa and therefore only examined a narrow slice of African cassava germplasm diversity (Rabbi et al., 2014b).

A limited genetic base for the dominant resistance implies potential vulnerability if the cassava mosaic geminivirus can evolve to overcome it. This possibility necessitates diversification of resistance sources to ensure durability. In order to determine with greater certainty whether there are additional sources of CMD resistance in the continent’s breeding germplasm, we undertook a large Genome-Wide Association Study (GWAS) using over six thousand cassava accessions from West and East Africa genotyped at more than 40,000 SNP loci using genotype-by-sequencing (GBS) approach (Elshire et al., 2011). The entire collection represents five sub-populations (Table 1) that are part of an ongoing international genomic selection-based breeding project in cassava (http://www.nextgencassava.orgl). In addition, we combined GWAS and genomic prediction in order to not only dissect the genetic architecture of resistance to CMD but also to assess the potential for population improvement by genomic selection (GS). We used a variety of approaches to localize and identify candidate genes for future investigation. The potential for GS to improve CMD resistance and for non-additive models to predict total genetic merit of clones for the selection of superior CMD resistant varieties were assessed. Finally, multi-kernel genomic prediction models were used to study the relative importance of qualitative and quantitative resistance sources.

**Table 1.**
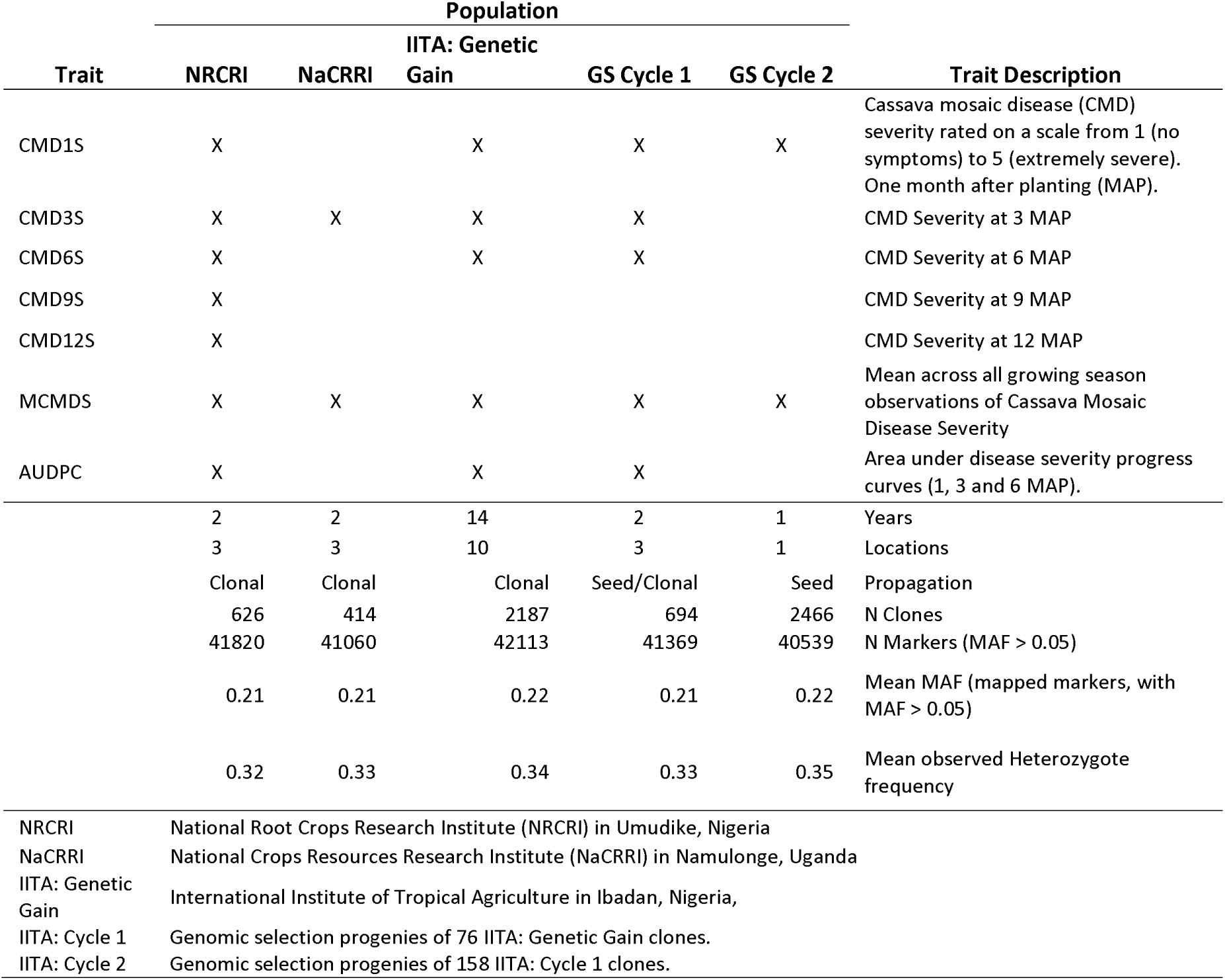
Summary of phenotype and genotype datasets analyzed.

## MATERIALS & METHODS

### Germplasm collection

The germplasm included in this study represent the reference populations used to develop genomic prediction models as part of a collaborative project between Cornell University and three breeding institutions: The International Institute of Tropical Agriculture (IITA) in Ibadan, Nigeria, the National Root Crops Research Institute (NRCRI) in Umudike, Nigeria and the National Crops Resources Research Institute (NaCRRI) in Namulonge, Uganda. The IITA’s Genetic Gain population is comprised of 694 historically important, mostly advanced breeding lines that have been selected and maintained clonally since 1970 (Okechukwu and Dixon, 2008; Ly et al., 2013). Most of these materials are derived from the cassava gene-pool from West Africa and early introductions of CMD tolerant parents derived from the inter-specific hybridization program at the Amani Station in Tanzania (Hahn et al., 1980b). It also includes hybrids of germplasm introduced from Latin America (see Table S1 for a list of accessions and details on pedigree where available). The NRCRI population contains 626 clones from their breeding program, 189 of which are also part of IITA’s Genetic Gain. The remainder of the NRCRI collection includes a large number of materials either directly from or derived with parentage from the International Center for Tropical Agriculture (CIAT) in Cali, Columbia (Table S2).

There are two major clades of cassava mosaic virus species, African Cassava Mosaic Virus (ACMV) and East African Cassava Mosaic Virus (EACMV) (Legg and Fauquet, 2004). EACMV is generally more severe in its symptoms and is present in west Africa but only in low proportion to ACMV, usually occurring as a dual infection (Legg and Fauquet, 2004; Rabbi et al., 2014b). This fact makes it all the more important to include east African cassava breeding germplasm in a more comprehensive screen of the genetic architecture of CMD resistance. The NaCRRI in Uganda has a population of 414 clones that represent the genetic diversity of East African cassava gene pool. The pedigree of this population arises from 49 parents coming from IITA, CIAT in Columbia and Amani Research Station in Tanzania (Table S3). The population was generated in part by making crosses of parents with qualitative resistance to parents with quantitative resistance as well as quantitative x quantitative and qualitative x qualitative resistances.

We also analyzed a large genotyped and phenotyped multi-parental population of individuals from two cycles of genomic selection (GS) conducted at IITA. The GS program at IITA will be described briefly here and in detail as part of other publications. In 2012 the IITA Genetic Gain population was used as the reference population from which genomic estimated breeding values (GEBVs) were obtained using the genomic BLUP method (GBLUP) (Heffner et al., 2009). Selection of clones from the Genetic Gain was based on a selection index including CMD and cassava bacterial blight disease severity and yield components (dry matter content, harvest index and fresh root yield). In the end, 83 parents gave rise to 2187 progenies, which we will call IITA Cycle 1. In 2013, the GEBVs for Cycle 1 were obtained, again using the Genetic Gain as a reference population and 84 Cycle 1 plus 13 (97 total) Genetic Gain clones were selected as parents, giving rise in 2014 to 2466 progenies (Cycle 2). The pedigrees of IITA Cycle 1 and Cycle 2 are available in Tables S4-S5.

### Phenotyping Trials

Phenotypic data were combined from trials conducted at multiple locations in Nigeria and Uganda. The data are contributed from all three breeding programs (IITA, NRCRI and NaCRRI). IITA’s Genetic Gain trials were conducted in seven locations over 14 years (2000 to 2014) in Nigeria. Each Genetic Gain trial comprises a randomized, unblocked design replicated one or two times per location and year. NRCRI’s population was phenotyped in two years, 2013 and 2014. During the 2012–2013 season the trial was conducted in one location, Umudike, Nigeria. In 2013–2014 the population was planted in three locations (Umudike, Kano, and Otobi). NRCRI’s trial design was a randomized incomplete block with three replications per location/year and 10 plants per plot. Trials at NaCRRI were conducted in two years: 2012-2013 and 2013-2014. In both years plots were 10 plants in two rows of 5 with randomized incomplete blocks. During the first year, a single location (Namulonge, Uganda) was used with only one replicate. During the second season, two replications were used at each of three locations: Namulonge, Kasese and Ngetta.

Genomic selection (GS) Cycle 1 (C1) progenies were observed as seedlings in the 2012-2013 field season with phenotyping conducted only for early disease expression and seedling vigor. Cycle 1 progenies were subsequently cloned and phenotyped in a three-location (Ibadan, Ikenne, and Mokwa) trial in 2013-2014 with all phenotypes scored. For the C1 clonal trial, planting material was only available for one plot of five stands per clone, so each clone was only planted in one of the three locations. Clones were assigned to each location so as to equally represent each family in every environment. The GS Cycle 2 (C2) individuals were observed in a seedling trial during the 2013-2014 field season. We note that expression of disease symptoms in cassava seedlings may not be representative of expression in clonal evaluations. This is in part because seedling symptoms can arise solely from whitefly transmission, making it probable that some asymptomatic plants are in fact escapes rather than resistant genotypes. Table 1 summarizes the phenotypes and phenotyping trials available for each sub-population. We also provide details about the sample sizes and replication numbers for each location/year of data analyzed (Table S6) and per accession (Table S7)

Cassava mosaic disease severity (CMDS) was scored on a scale of 1 to 5, with 1 representing no symptoms and 5 indicating the most severe symptoms. CMDS was scored at up to five time points (1, 3, 6, 9 and 12 months after planting) depending on the trial. Additionally, we analyze the season-wide mean CMDS score (MCMDS), which is used for making selections and the area under the disease progress curve (see below; AUDPC). The distribution of raw phenotypic data used in each population and for each trait can be seen in Figures S1–S6.

### Statistical Models and Analyses of Phenotypes

Our interest in this study was to identify key aspects of the genetic architecture of cassava in Africa rather than location- or year-specific QTLs. We condensed up to 38854 observations on 6198 genotyped and phenotyped clones to single BLUPs for each. To do this, we fit the following mixed linear model with the *lme4* package in R:

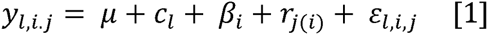

Here, y_l,i,j_ represents raw phenotypic observations, μ is the grand mean, *C_l_* is a random effects term for clone with 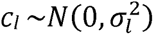, β_i_ is a fixed effect for the combination of location and year harvested, r_j(i)_ is a random effect for replication nested within location-year combination assumed to be distributed 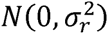 and finally, ε_l,i.j_ is the residual variance, assumed to be random and distributed *N*(0, *a*^2^). Because the number of observations per clone varies greatly in our dataset (from 1 to 941, median of 2; Table S7), we expect BLUPs are differentially shrunken to the mean. To counter this, we de-regressed BLUPs according to the following formula:

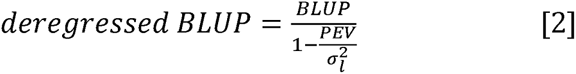

Where PEV is the prediction error variance for each clone and 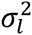 is the clonal variance component. The distribution of deregressed BLUPs used as response variables in GWAS can be seen in Figures S7-S13.

We also calculated areas under disease progress curves (AUDPC) for each clone using data from 1, 3 and 6 months after planting. To do this, we treated severity scores from any time point as the same trait with a second variable indicating the time point of the score. We then ran the model indicated in [1] but with C_l_ indicating the clone-time point combination. This gave us a deregressed BLUP for each clone at each time point. We calculated areas under these curves using the trapezoid rule as implemented by the *auc* function in the *flux* R package (http://cran.r-proiect.org/web/packages/flux/index.html). We excluded 9 and 12 months data because they were only scored at NRCRI, thus including them would have limited the ability to compare results for this trait between populations.

### Genotype Data

Genotyping of SNP markers was done by the genotyping-by-sequencing procedure (Elshire et al., 2011) using the ApeKI restriction enzyme recommended by (Hamblin and Rabbi, 2014) and read lengths of 100bp. Marker genotypes were called with the TASSEL GBS pipeline V4(Glaubitz et al., 2014) and aligned to the cassava version 5 reference genome, available on Phytozome (http://phvtozome.jgi.doe.gov) and described by the International Cassava Genetic Map Consortium (2014). Individuals with >80% missing SNP calls and markers with more than 60% missing were removed. Also, markers with extreme deviation from Hardy-Weinberg equilibrium (Chi-square >20) were removed. Allele calls were maintained if depth was ≥2 otherwise the call was set to missing. Marker data was converted to dosage format (0,1, 2) and missing data were imputed with the glmnet algorithm in R (http://cran.r-project.org/web/packages/glmnet/index.html) as described in(Wong et al., 2014). In order to judge the resolution of association analyses we calculated pair-wise linkage disequilibrium (LD) between all markers with a MAF of 5% on each chromosome using PLINK (version 1.9, https://www.cog-genomics.org/plink2). We examine the rate of decay with increasing physical distance between markers.

### Population Structure and Genome-Wide Association Analyses

In order to examine the patterns of relatedness within and among our populations and to control population structure, we constructed a genomic-relationship matrix according to the formulation of Van Raden (VanRaden, 2008); see also 25), as implemented in PLINK, using all markers with greater than 1% minor allele frequency (MAF). We also use this relationship matrix for genomic prediction (see below).

We conducted principal components analysis (PCA) on SNP markers with MAF > 5% using the *prcomp* function in R. PCA on SNP markers is often used to identify major patterns of relatedness (population structure) in a sample and the first few PCs can be used as covariates to control false-positive rates in GWAS(Price et al., 2006). Because the genomic selection progenies (C1 and C2) are by far the largest part of our dataset and because these individuals are descended from the IITA Genetic Gain population, we excluded these from the initial PCA. We then projected these individuals into the genetic space defined by the three training populations (NRCRI, IITA, NaCRRI) using the *predict* function in R. This allows us to visualize and quantify the relatedness in our populations based on the founders only and unbiased by the large size of the C1 and C2 collections.

Because GWAS has not previously been done in this or any other cassava collection, we tested several different models for controlling population structure. In particular, we compared the genome-wide inflation of p-values between a general linear model (GLM) with no population structure controls, a GLM with 5 principal components (GLM + 5 PCs)(Price et al., 2006), and a mixed linear model (MLM), which fits a random effect for clone with ~N(0,σ_g_^2^K), where σ_g_^2^ is the clonal variance component and K is the relationship matrix described above (Kang et al., 2010). MLM were conducted using the P3D and compression method (Zhang et al., 2010). All GWAS were conducted in TASSEL (version 5, [27]). We compare the observed – log_10_(p-values) against the expectation using QQ-plots. We used visual inspection of QQ-plots to judge which model most effectively reduced the genome-wide inflation of −log_10_(p-values) typically attributed with population structure. We consider association tests significant when more extreme than the Bonferroni threshold (with experiment-wise type I error rate of 0.05).

Because marker effects, LD patterns, and allele frequencies may differ within as well as across sub-populations, we conducted GWAS population-wide as well as within each sub-population. In each analysis, we used markers that segregated with MAF > 5% in that specific sub-population. Bonferroni thresholds were calculated according to the number of markers analyzed in each sub-population.

We also examined the proportion of variance in the deregressed BLUPs explained by the kinship matrix, K using the variance components estimated when TASSEL fits the MLM model.

### Candidate Genes

Because the underlying mechanisms of plant disease resistance are of general interest and identification of causal polymorphism may aid in transgenic approaches and/or marker assisted selection, we identified candidate genes in CMD associated regions. Significant SNPs from the GWAS results corresponding to four time points (1, 3, 6 and 9 months after planting) were selected for the analysis. We considered SNPs that were both above the Bonferroni threshold and were located within exons or introns of cassava genes. The SNP position on the genome was compared with the gene interval position using the annotation list from Phytozome 10. Gene ontology annotation for each time point and combining all the datasets was done with Panther (http://go.pantherdb.org/). We have generated whole genome sequences (WGS) from one CMD resistant clone (I011412) and two CMD tolerant clones (I30572 and TMEB1). TMEB1 is a landrace from Ogun State, Nigeria also called Antiota, that is not likely to contain the qualitative resistance allele and is usually classified as tolerant or only partially susceptible to CMD (Raji et al., 2008; Rabbi et al., 2014b). Similarly, 130572 is an improved variety whose parents were the *M. glaziovii*-derived clone 58308 and a south American Cassava (Branca de Santa Catarina) and is therefore known to have only the quantitative resistance source (Fauquet et al., 1990). *PCR*-free libraries were generated from these clones and sequenced at 20X coverage using Illumina HiSeq. Additionally two resistant clones (TMEB3 and TMEB7) were obtained from Phytozome (http://phytozome.jgi.doe.gov). TMEB3 is itself the original landrace parent from which the qualitative resistance source has been derived and TMEB7 has been shown to be nearly genetically identical to TMEB3. We therefore define TMEB3, TMEB7 and I011412 as “resistant” lines while TMEB1 and I30572 will be referred to, for simplicity as “susceptible” primarily on the basis of whether they do or do not have the qualitative resistance source *CMD2.* These sequences were aligned against the cassava V5 reference genome assembly to call the variants to identify the genomic difference between resistance and susceptible clones in candidate gene loci. Since the genotypes compared were few in number, we called SNPs manually using an exon annotated sequence and the Integrative Genomics Viewer software (IGV; http://www.broadinstitute.org/igv/).

### Genomic Prediction of Additive and Total Genetic Merit

We used a multi-random effects (a.k.a. multi-kernel or multi-relationship matrix) genomic prediction model to compare the variance explained and prediction accuracy achieved from the major CMD QTL (CMD2) compared to the rest of the genome. Specifically, we created relationship matrices either from all markers, markers significantly associated with CMD2 from GWAS results, or all markers *not* in the region of the QTL.

For additive relationships, we used the formulation described above for controlling population-structure (VanRaden, 2008). Dominance relationships can be captured as 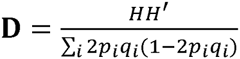 (Su et al., 2012; Muñoz et al., 2014). Where H is the SNP marker matrix (individuals on rows, markers along columns), heterozygotes are given as (1 - 2p_i_q_i_) and homozygotes are (0 - 2p_i_q_i_). We made a custom modification (available upon request) to the *A.mat* function in the *rrBLUP* package (Endelman, 2011) to produce the **D** matrix. Relationship matrices that capture epistasis can also be calculated by taking the hadamard product (element-by-element multiplication; denoted #) of two or more matrices (Henderson, 1985). For simplicity, we tested an additive-by-dominance **(A#D)** matrix in this study.

We tested four models. Model 1 used all markers and only a single, additive kernel (Additive_All_Markers_). Model 2 used all markers but three kernels, Additive_All_Markers_ + Dominance_All_Markers_ + Epistasis_All_Markers_. Model 3 used two additive kernels, one constructed from the 163 CMD2 significant markers (Additive_CMD2_) and the other from all markers outside of the chromosomal region bounded by CMD2 markers (Additive_Non-CMD2_). Model 4 had four kernels: Additive_CMD2_ + Dominance_CMD2_ + Epistasis_CMD2_ + Additive_Non-CMD2_.

We assessed the influence that modeling non-additive genetic variance components have on genomic prediction using a cross-validation strategy (see below). We used the deregressed BLUPs for MCMDS as described above. In our data, the number of observations per clone ranges from one to 941 (checks, TMEB1 and I30572) with median of two and mean of 10.6 (Table S7). Pooling information from multiple years and locations, especially when there is so much variation in numbers of observations can introduce bias. Much theoretical development, particularly in animal breeding has been done to address this issue, and we followed the approach recommended by Garrick et al. (2009)

In the second step of analysis, where deregressed BLUPs are used as response variables, weights are applied to the diagonal of the error variance-covariance matrix **R.** Weights are calculated as 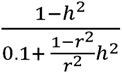, where *h^2^* is the proportion of the total variance explained by the clonal variance component, 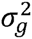 derived in the first step **(Garrick et al. 2009).**

We implemented a 5-fold cross-validation scheme replicated 25 times to test the accuracy of genomic prediction using the genomic relationship matrices and models described above. In each replication, we randomly assign each individual to one of five groups (folds). We then select one fold, remove the corresponding individuals from the training set and use the remaining four folds to predict the fold that was left out. We iterate this process over each of the five folds to produce a prediction for each individual that was made while its phenotypes were unobserved. For each replicate of each model, we calculated accuracy as the Pearson correlation between the genomic prediction made when phenotypes were excluded from the training sample and the BLUP (ĝ, not-deregressed) from the first step. For each model, we calculated accuracy both of the prediction from the additive kernel (where present) and the total genetic merit prediction, defined as the sum of the predictions from all available kernels (e.g. additive + dominance + epistasis). Genomic predictions were made using the *EMMREML* R package. For simplicity, we tested only the trait MCMDS in the IITA Genetic Gain population.

In addition, we assessed the predictability of CMD based on random forest regression (RF), a non-linear, machine-learning approach that excels at capturing non-additive especially interaction-type genetic effects(Jannink et al., 2010). We used RF only with the significant CMD2 associated markers as predictors to assess additional evidence for interaction at this locus on the basis of prediction accuracy achieved. We used the same cross-validation scheme described above.

## RESULTS

### Genotyping Data

SNP marker data was generated using genotyping-by-sequencing (GBS) (Elshire et al., 2011). Overall coverage was 0.07x (range 0.05-0.2). There were 114,922 markers that passed initial filters with a MAF > 1%. Of these, 95,047 are mapped to the genome. Of mapped markers used for GWAS (MAF > 5%), there was an average of 2293 SNPs per chromosome or one marker every 9.5 kb. The mean MAF (0.21-0.22), mean heterozygosity (0.32-0.35) and number of markers analyzed (40,539-42,113) were similar between sub-populations (Table 1). Most chromosomes in most populations had mean r^2^ > 0.2 extending 10 to 50 Kb. The r^2^ between markers 4.5-5.5kb apart was 0.3 on average (median 0.13) suggesting at least some LD between most causals and at least one marker but also that increased density in future studies will provide additional mapping resolution (Figs S14-S19).

### Population stratification and structure

Principal components analysis of our SNP dataset revealed subtle differentiation among African cassava clones analyzed. This can been seen from a plot of the first four PCs (cumulative variance explained = 15%). The Nigerian sub-populations (NRCRI, IITA Genetic Gain, Cycle 1 and Cycle 2) occupy similar genetic space, but the Ugandan sub-population (NaCRRI) is somewhat distinct on PC1 and PC2. This may be consistent with a history of germplasm sharing and recurrent use of elite parents among African breeding institutes.

We tested several standard GWAS models for controlling inflation of p-values caused by population structure including a general linear model (GLM, no correction); a GLM with the first 5 PCs of the SNP matrix as covariates; and a mixed-linear model using the marker-estimated kinship matrix. Visual inspection of QQ plots (Fig. 2 inset, Figs. S20-25) indicated that the MLM was most consistent for reducing −log_10_(p-values) towards the expected level (i.e. controlling false-positives, removing population structure effects). All subsequent results are therefore based on mixed-model associations. From the variance components estimated when fitting MLMs we found that kinship matrices explained on average 57% (range 31-94%) of the phenotypic variance (Table S1).

**Figure 1.**
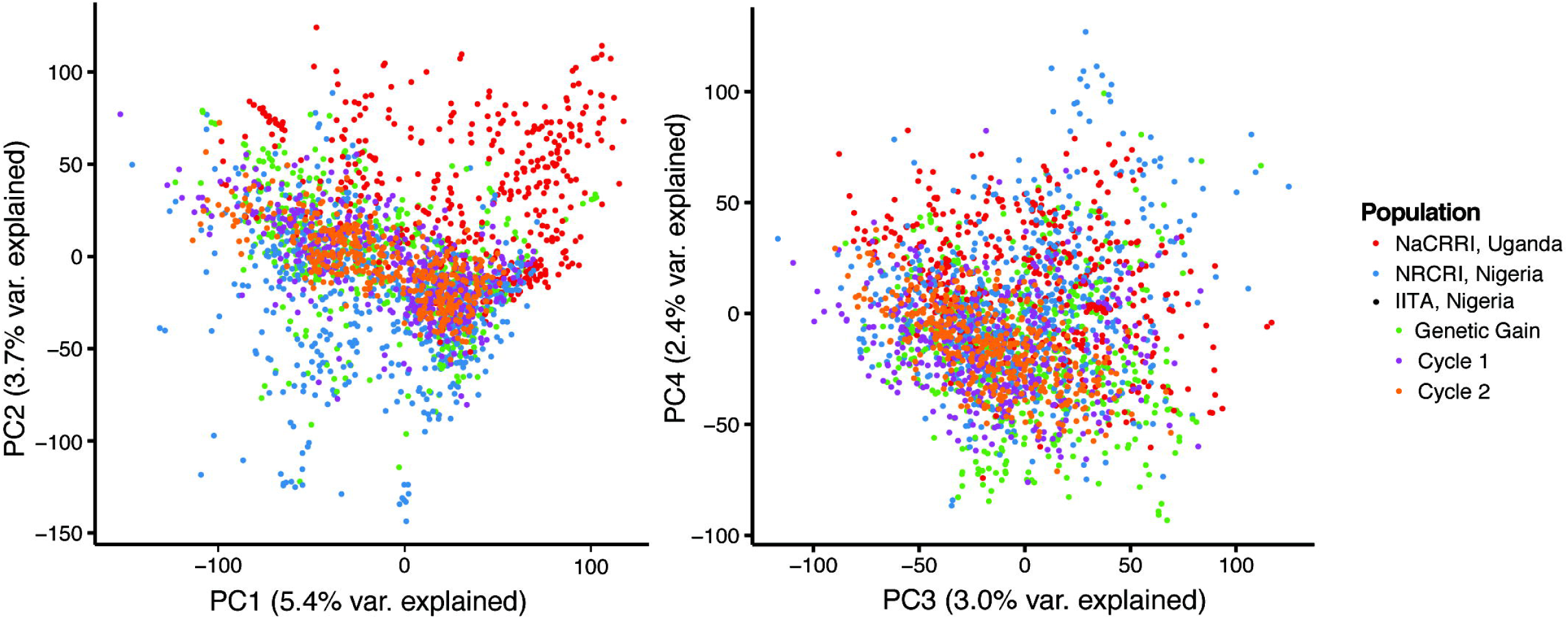
Plot of the first four principal components of the SNP marker matrix. The three main training populations were used in the PCA and are shown here. A random sample of IITA GS Cycles 1 and 2 were projected into the genetic space and are displayed here.

### Overall Genome-wide Associations

Association tests were performed for CMD symptom severity at one, three, six, nine and twelve months after planting (where measured) in the five sub-populations (Table 1) and in analysis that combined all accessions. We identified 311 markers in total that pass a Bonferroni significance threshold (Fig. 2, Table S8). However, many significant SNPs were detected because of rare marker genotypes that were phenotypically extreme (Figure S26). The F-test implemented by TASSEL is sensitive to imbalanced sample size between groups and we wish to be conservative and only consider significant results that we can be confident in. Therefore, we only consider SNPs where each genotype class (e.g. aa, Aa, AA) is represented by at least 10 individuals. This reduced the number of significant markers to 198, on 14 chromosomes, mostly concentrated at a single region of chromosome 8. Significant results were found within each sub-population, with more signals associated with greater sample size (e.g. Cycle 1). Variance explained by significant markers ranged from 0.5% to 22% (median 3.5%) (Table S9).

**Figure 2.**
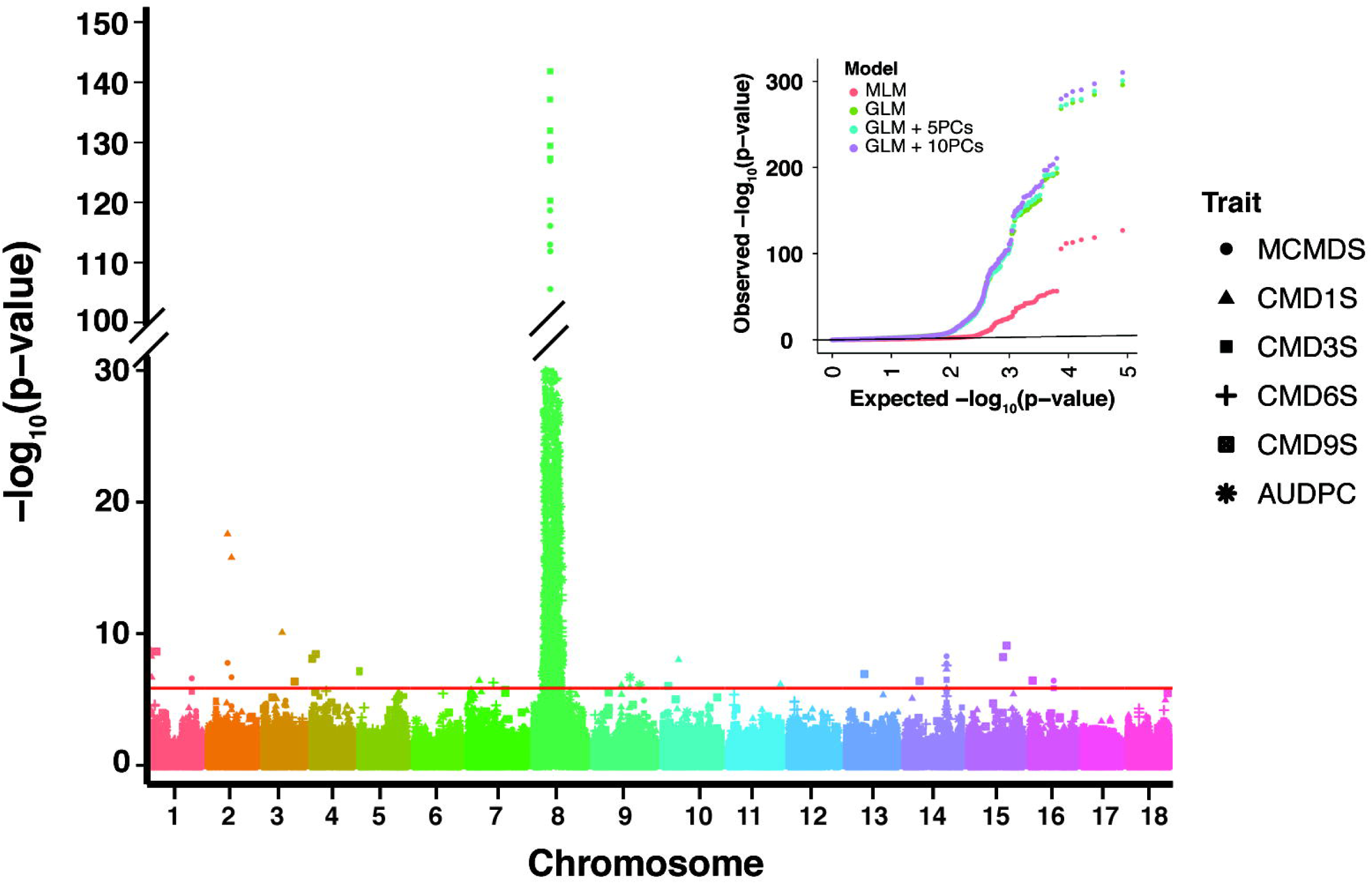
Manhattan plot from mixed-linear models summarizing genome-wide association results for all traits in all sub-populations. Bonferroni significance threshold is shown in red. An example QQ-plot (MCMDS in the population-wide analysis) is shown inset to demonstrate the differences between various population structure controls.

### Chromosome 8 contains the major resistance locus, *CMD2*

There were 163 significant markers on chromosome 8 (between 3.56-11.38 megabases; Fig. 3a) with the top marker (S8_7762525) explaining 5-22% of the variance depending on the sub-population. The frequency of the resistance-associated allele at S8_7762525 is 56% overall (range: 44% in IITA Genetic Gain to 63% in IITA Cycle 2 progenies).

**Figure 3.**
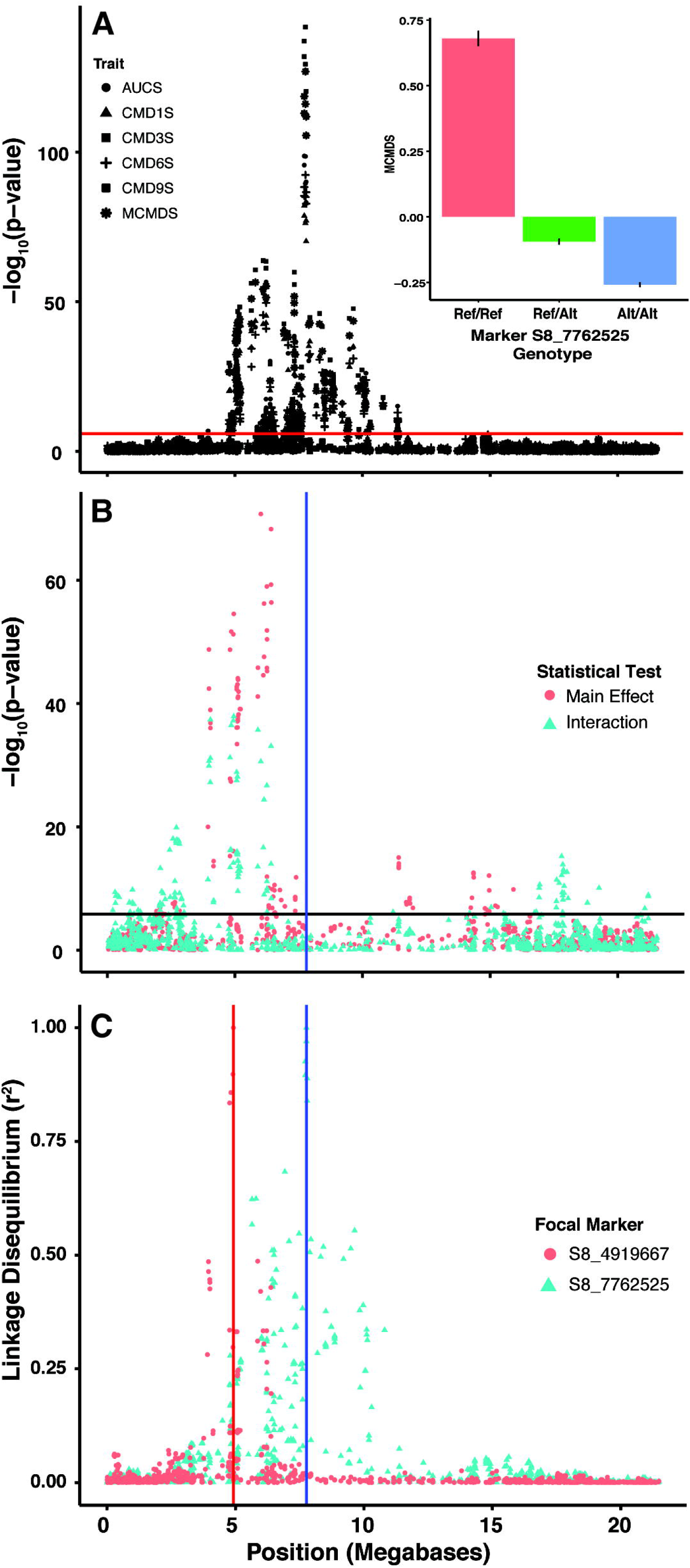
Plots dissecting the major effect QTL on chromosome 8. (A) Manhattan plot summarizing genome-wide association results for all CMD-related traits in the population-wide mixed-linear model analysis, zoomed to chromosome 8 only. (B) Manhattan plot showing linear model tests for interactions (blue dots) between the top marker (S8_7762525, blue vertical line) and every other maker on chromosome 8. Red dots are for the main effect of the second marker (main effects of S8_7762525 are not shown). (C) LD between S8_7762525 (blue vertical line) and every other maker (blue dots), plus LD between the marker with the strongest interaction effect (S8_4919667; red vertical line) and every other marker. Bar plot showing mean and standard error for MCMDS between each genotype class at the top marker, S8_7762525 (Inset).

The resistance allele at S8_7762525 is incompletely dominant (Fig. 3 inset); homozygotes with the alternate allele were closer to CMD free than heterozygotes. To formally test this, we conducted a post-hoc test for an additive effect at this marker that explained 15% of the variance compared to a test of additive plus dominance effect that explained 20%, and a test of dominance alone that accounted for only 2%.

We confirmed that our major QTL is the *CMD2* locus by aligning previously published SSR marker primers (SSRY28, NS158 and SSRY106) (Akano et al., 2002; Lokko et al., 2005; Okogbenin et al., 2007, 2012a; Mohan et al., 2013) to the reference genome using E-PCR (http://www.ncbi.nlm.nih.gov/tools/epcr/). Our significant markers on chromosome 8 co-locate with these markers (Fig. S28a). Additionally, scaffolds bearing the significant QTL reported in Rabbi et al. (2014a; b) are located in this region. However, while Rabbi et al.’s (2014a; b) strongest association was on scaffold 5214, corresponding to Chr. 8 position 6511133, the strongest association for the present study is on scaffold 6906 (7454373-7836749), more than a megabase away. This discrepancy is due to the fact that the SNP markers in scaffold 6906 did not segregate in the resistant parents of the bi-parental mapping populations.

### Dissecting resistance originating from alleles or loci on Chromosome 8

The significance region on chromosome 8 is large (~8 Mb; Fig. 3a). In fact, the region appears as two, sometimes equally significant peaks in some sub-populations (Fig. S27). We scanned the region for haplotype blocks with PLINK (version 1.9, https://www.cog-genomics.org/plink2) and found it was not characterized by a single, or even a few large, but many small LD blocks (Fig. S29). A second locus (*CMD3*) has been reported on the same chromosome as *CMD2*(Okogbenin et al., 2012b). The authors reported the marker NS198 to be 36 cM from *CMD2* and associated with very strong resistance in the progeny of TMS972205. E-PCR collocated NS198 on chromosome 8, five megabases (position 997099) outside our significance region (Fig. S28). Thus our results suggest a second QTL (i.e. *CMD3*), if present, is much closer to *CMD2* than previously believed.

We used several approaches to evaluate the evidence for multiple QTL in the region. We conducted a post-hoc test for interactions between the top-marker on chromosome 8 and every other marker on the chromosome. There were significant interactions, explaining up to 42% of the variance, 1-3 megabases from the top GWAS hits, but none in the region surrounding S8_7762525 (Fig. 3b, Fig. S30).

We implemented a multi-locus mixed-model (MLMM (Segura et al., 2012)), which uses a forward-backward stepwise model selection approach to determine which and how many marker cofactors are required to explain the associated variance in the region. The MLMM for MCMDS in the population-wide sample selected five markers (S8_7762525, S8_6380064, S8_6632472, S8_7325389, S8_4919667) spanning the significance region (Fig. S31). Of the five, the first was our top marker S8_7762525, the fourth is only about 400 Kb away, and the remaining three cover the secondary peak and the region of statistical interaction. These markers are mostly in linkage equilibrium (Table S10) and collectively explain up to 40% of the variance. The selection of markers distributed across the region by MLMM including both putative peaks to explain the phenotypic-association in the region is additional evidence in support of a multi-locus hypothesis.

LD decays in the region to low levels (r^2^<0.25) and is virtually zero between significant markers on the left, e.g. S8_5064191 and those on the right, e.g. S8_762525 of the significance region (Fig. 3c). Combined with our genome-wide analysis of LD decay rates (Figs. S14-S19) this LD decay makes it unlikely that a single locus or allele is responsible for the associated region.

We examined the two-locus genotype effects (e.g. between S8_7762525 and the SNP with the most significant interaction test, S8_4919667) and found a usually dominant effect of the secondary resistance allele (e.g. S8_4919667) in the heterozygous and homozygous resistant background at the primary peak (Fig. 4; Fig. S32). We found little evidence of secondary peak effects in the homozygous susceptible primary peak background. Clones that are homozygous resistant at both loci are superior to all other cassava clones, expressing virtually no symptoms (Fig. 4c).

**Figure 4.**
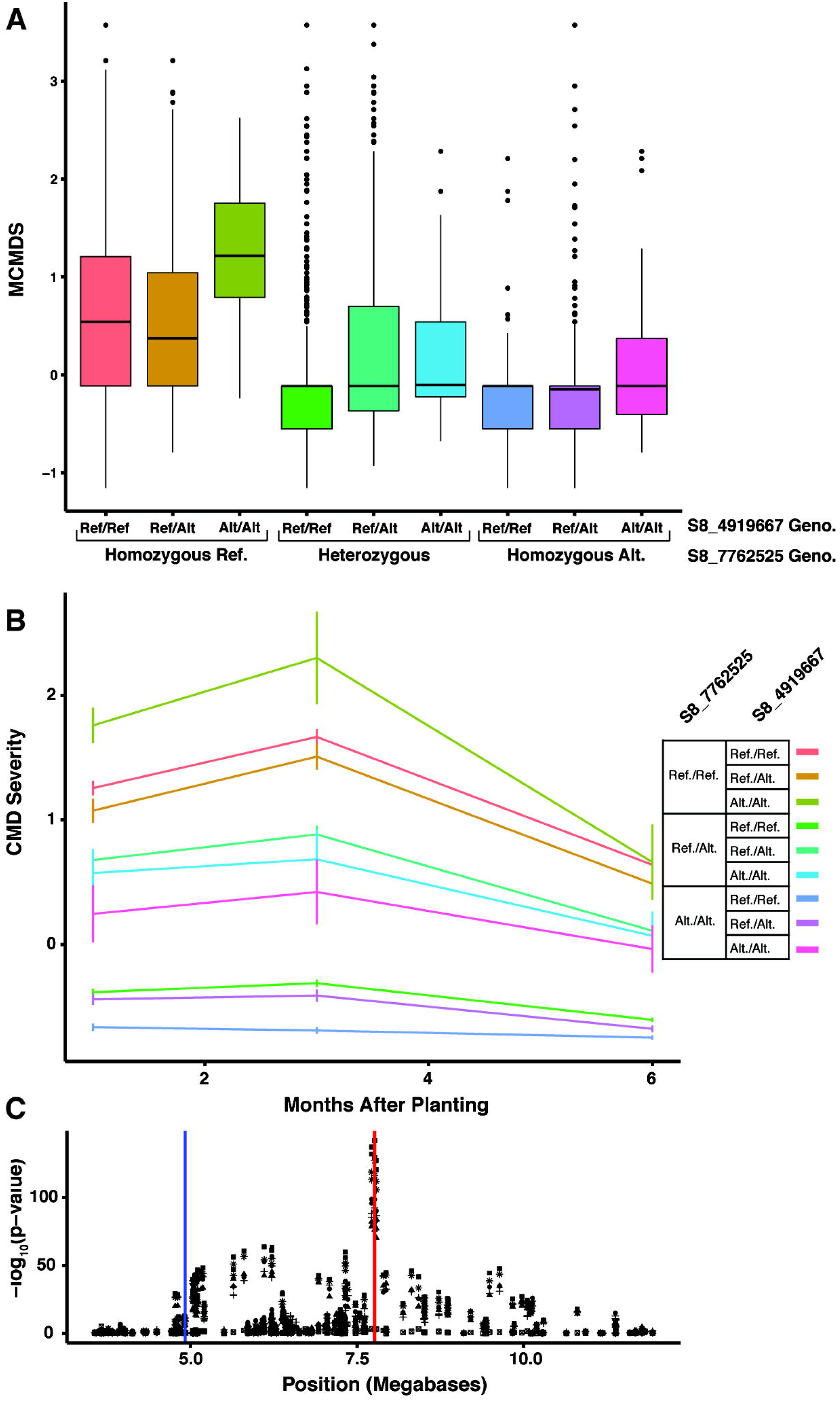
Plots demonstrating the combined effect of the genotype at the top marker (S8_7762525) and the most epistatic marker (S8_4919667). (A) Boxplot showing the distribution of mean CMD severity scores (MCMDS) for each two-locus genotype. (B) Disease progress curves showing mean and standard error CMD severity across 1, 3 and 6 months after planting for each two-locus genotype. (C) Zoomed Manhattan plot showing the location of the two markers being compared; S8_7762525 (red line) and S8_4919667 (blue line).

### Other Loci Associated with CMD Resistance

We identified thirty-five markers on 13 chromosomes that explained 0.5–10% (median 4%) of the variance (Table S9). Many of these had recessive and usually rare susceptibility alleles (Fig. S26). Marker S4_637212 explained 4% of the variance (CMD6S, Genetic Gain) and had an additive effect. Marker S11_20888811’s recessive resistance allele appears to lower CMD symptoms 4% more than *CMD2* (S8_7762625) but only 14 clones are homozygous resistant at this locus (Fig. S26). Further work on this locus is urgently needed to determine it’s possible impact as its frequency increases. There were four significant markers on chromosome 14 with mostly dominant effects and explaining up to 5% of the variance. Two previously published SSR markers (SSRY44, NS146) (Mohan et al., 2013) are located within 1.4 megabases of these SNPs (Fig. S28b). Four markers, spread across seven megabases of chromosome 9, with recessive susceptibility loci, explained up to 10% (S9_14551208) of the variance. These markers co-located with SSRY40, originally reported as *CMD1* and associated with quantitative resistance (Fregene et al., 2000; Mohan et al., 2013).

### Candidate Genes

We intersected our association-results with available gene annotations and related data and identified 105 unique genes within the association peaks, with 79, 61, 56 and 9 genes identified at one, three, six and nine months after planting, respectively (Table S11, Fig. S33). There were no significant differences between gene ontology categories between time points. Most of the annotated genes are involved in metabolic processes (Fig. S34). Thirty-five out of the 105 genes are known to respond to cassava mosaic virus infection (Allie et al., 2014) (Table S11).

Among these genes we found ones known to be susceptibility or resistance factors, a number of which are also involved in plant-geminivirus interaction processes (Hanley-Bowdoin et al., 2013). We found two peroxidases Cassava4.1_029175 and Cassava4.1_011768 within the primary QTL region (scaffold 6906, ~7.7Mb); peroxidases are pathogenesis-related proteins (PRs), involved in host response to infection(van Loon et al., 2006). In the secondary GWAS peak (scaffold 5214, 5-6Mb) six SNPs were in a protein disulfide-isomerase like 2-2 ortholog, a thioredoxin (PDIL2-2, cassava4.1_007986). In barley, an ortholog of PDIL2-2 (*HvPDIL5-1*) is a known virus susceptibility factor as are *PDI* gene family members across the animal and plant kingdoms (Yang et al., 2014). We also identified the Ubiquitin-conjugating enzyme E2 ortholog (UBC5) gene (cassava4.1_017202) under the secondary GWAS peak (scaffold 5214, 5-6 Mb region). Genes like UBC5 in the ubiquitinylation pathway have been known to influence plant virus infection response (Becker et al., 1993).

We analyzed the coding sequence of the three genes mentioned above in three CMD resistant cassava genotypes known to possess the qualitative resistance allele(s) (TMEB3, TMEB7 and I011412) and in two susceptible/tolerant ones known to possess only quantitative resistance sources (I30572 and TMEB1). We identified SNPs within the coding regions and identified amino acid changes (Table S12). Two non-synonymous mutations were found on exons 7 and 9 of Cassava4.1_007986, homozygous in the susceptible group but heterozygous in the resistant clones (Table S12). The peroxidase, Cassava4.1_011768 did not show any non-synonymous mutations specific to the resistant/susceptible group. However, Cassava4.1_02917, showed three non-synonymous mutations that were specific to the susceptible group.

### Genomic Prediction of Additive and Total Genetic Merit

We tested four prediction models using cross-validation: (1) Additive_All_Markers_, (2) Additive_All_Markers_ + Dominance_All_Markers_ + Epistasis_All_Markers_, (3) Additive_CMD2_ + Additive_Non-CMD2_, (4) Additive_CMD2_ + Dominance_CMD2_ + Epistasis_CMD2_ + Additive_Non-CMD2_. Mean cross-validation accuracy averaged 0.53 for additive and 0.55 for total value across models (Table 2, Figure 5). Including non-additive effects, using all markers (model 2) shifted 60% of the variance to dominance and epistasis and decreased the accuracy of the additive prediction from 0.53 (model 1) to 0.51, but gave increased total prediction accuracy of 0.55. An additive only model giving separate weight to CMD2 and non-CMD2 regions (model 3) had the highest total prediction accuracy (0.58), with most accuracy coming from CMD2 (0.54) vs. non-CMD2 (0.29) but most variance absorbed by non-CMD regions. Modifying model 3 to allow the CMD2 region additive, dominance and epistatic effects (model 4) slightly decreased total prediction accuracy (0.57) relative to model 3, with most accuracy coming from the additive CMD2 kernel (0.52), but with 51.7% non-additive variance, 33.6% non-CMD2 variance and only 14.7% additive CMD2.

**Table 2.**
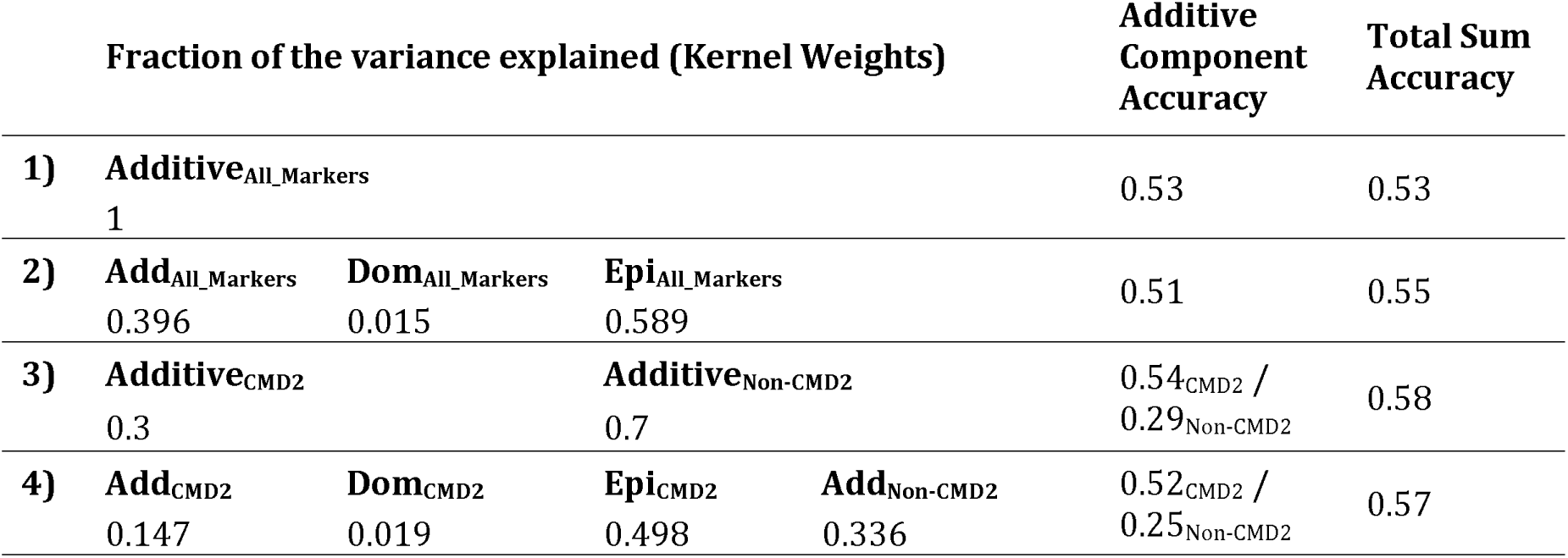
Summary of cross-validation results for MCMDS. Mean kernel weights as well as additive and total prediction accuracies are reported for each of four models tested.

**Figure 5.**
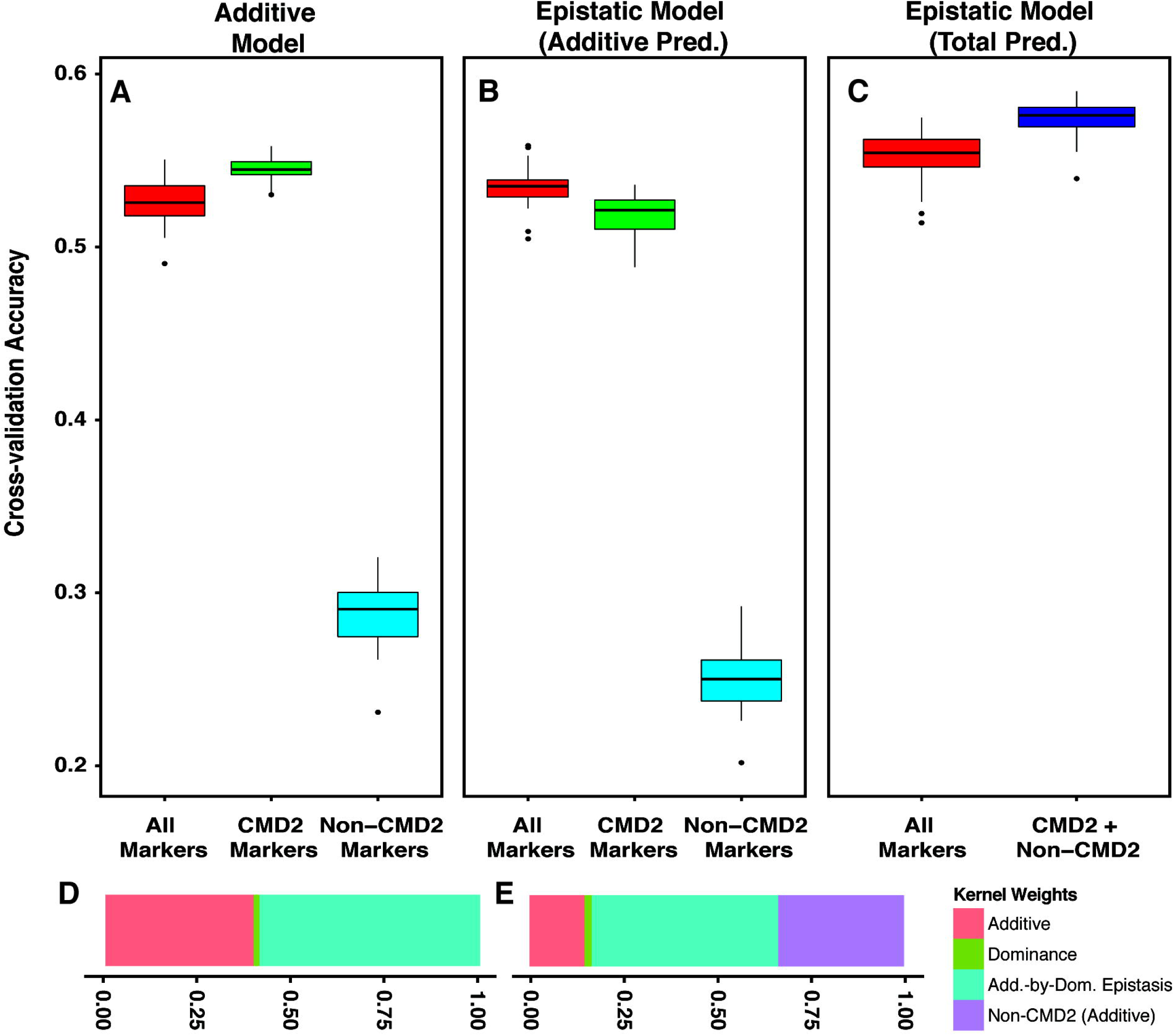
Cross-validated genomic prediction results for MCMDS. (A-C) Box plots of accuracies from 25 reps of 5-fold cross-validation. (A) Accuracies of the additive models (#1 and #3) using either a single kernel (Additive_All_Markers_) or two-kernels (Additive_CMD2_ and Additive_Non-CMD2_). (B) Accuracies of the additive predictions from the models that included dominance and additive-by-dominance epistasis (models #2 and #4). A single additive accuracy is calculated from the model #2 (Additive_All_Markers_) and two accuracies for model #4 (Additive_CMD2_ and Additive_Non-CMD2_). The accuracy of total genetic merit prediction from models (#2, all markers; #4, CMD2 + Non-CMD2) with dominance and epistasis are shown in (C). Kernel weights corresponding to the partitioning of the genetic variance for the epistatic models are shown in (D, all markers model #2) and (E, CMD2 + Non-CMD2 model #4).

## DISCUSSION

The present study solidifies our understanding of the genetic resistance to CMD that is available in African cassava germplasm and demonstrates the efficacy of genomic selection at improving CMD resistance. After conducting the first genome-wide association study for this species with markers anchored to chromosomes, we are able to confirm that the basis of genetic resistance to CMD is indeed narrow, arising chiefly from a single region of chromosome 8 that collocates with the loci *CMD2* (Akano et al., 2002) and *CMD3* (Okogbenin et al., 2012b). The lack of new major effect loci is a key outcome of our study. Even after analyzing a broad sample of the breeding germplasm from West and East Africa. However, we also identified 13 regions of small effect including one on chromosome 9 that collocates with *CMD1* (Fregene et al., 2000).

Another key result of our analysis is that the most highly resistant cassava clones, those that never show disease symptoms, are only identified using models of epistasis in the significance region on chromosome 8. We propose two alternative hypotheses to explain this result. As suggested both in our analyses and previous studies (Okogbenin et al., 2012b) there may be multiple interacting loci in the region (i.e. *CMD2* and *CMD3*). Alternatively, our results may arise from a complex haplotype structure, where observed levels of resistance come from a single locus with one moderate and another strong resistance allele segregating in the population. An example of the later scenario is resistance to tomato yellow leaf curl which initially mapped to two genes Ty-1 and Ty-3 on the same chromosome, but was later revealed by fine-mapping to be one gene with multiple alleles (Verlaan et al., 2013).

In order to facilitate functional studies of the qualitative resistance source(s) on chromosome 8, we used our GWAS results to identify three candidate genes. Interestingly, there are no major resistance genes (e.g. NBS-LRR) in our region of interest (Lozano et al., 2015). We found two peroxidases, which have recently been shown to down-regulate in response to cassava mosaic geminivirus infection in susceptible genotypes (Allie et al., 2014) and a thioredoxin, which can be important for plant defense activation (Bashandy et al., 2010; Ballaré, 2014). We note that our genome assembly contains gaps ((ICGMC), 2014) and is based on a South American accession (Prochnik et al., 2012) that may not possess the causal gene(s). Significant work remains to identify the causal mechanism of qualitative resistance to CMD.

Finally, we demonstrate the potential of genomic selection for CMD resistance breeding. In agreement with our association analyses, we found most of the variance and the prediction accuracy was attributable to the chromosome 8 QTL(s). While additive models will allow us to accurately select parents for cassava breeding, we found non-additive prediction of total genetic merit to be even more accurate. Prediction of total genetic merit will therefore enable cassava breeders to more easily identify clones with superior disease resistance to be elite varieties, effectively exploiting dominance and epistasis for crop improvement. Further, it is significant that, while accuracy is low for the quantitative (non-major gene) components, it is not zero. Thus it should be possible to do genomic selection to simultaneously improve both qualitative (i.e. *CMD2/CMD3*) and quantitative (i.e. polygenic background) resistance.

The results we present in this study will represent progress towards discovering the mechanistic basis for major gene resistance to CMD and will also aid breeders seeking to pyramid useful alleles and achieve symptom-free cassava varieties either by marker assisted or genomic selection. In only two years we have conducted two rounds of selection and recombination, twice as fast as conventional phenotypic selection, and have increased the resistance-allele frequency at our top marker from 44% to 63%. The present study is an example of the possibilities for rapidly improving and dynamically breeding a crop that is crucial for hundreds of millions, particularly in underdeveloped regions of the world.

## ACKNOWELDGEMENTS

We acknowledge the Bill & Melinda Gates Foundation and UKaid (Grant 1048542; http://www.gatesfoundation.org) and support from the CGIAR Research Program on Roots, Tubers and Bananas (http://www.rtb.cgiar.org). We give special thanks to A. G. O. Dixon for his development of many of the breeding lines and historical data we analyzed. Thanks also to A. I. Smith and technical teams at IITA, NRCRI and NaCRRI for collection of phenotypic data and to A. Agbona, A. Ogbonna, E. Uba and R. Mukisa for data curation. This work is dedicated to the memory of Martha Hamblin.

Abbreviations:

Genome-wide association analysis (GWAS); genomic selection (GS); Cassava mosaic disease (CMD); genotype-by-sequencing (GBS); International Institute of Tropical Agriculture (IITA); National Root Crops Research Institute (NRCRI); National Crops Resources Research Institute (NaCRRI); African Cassava Mosaic Virus (ACMV); East African Cassava Mosaic Virus (EACMV); genomic estimated breeding values (GEBVs); Cassava mosaic disease severity (CMDS)

